# Abolishing storage lipids induces protein misfolding and stress responses in *Yarrowia lipolytica*

**DOI:** 10.1101/2023.05.02.539027

**Authors:** Simone Zaghen, Oliver Konzock, Jing Fu, Eduard J Kerkhoven

**Affiliations:** Division of Systems and Synthetic Biology, Department of Life Sciences, Chalmers University of Technology, Göteborg, Sweden

**Author notes:** These authors contributed equally (Simone Zaghen, Oliver Konzock).

**Keywords:** Q4 strain, fat free *Yarrowia lipolytica*, lipid *Yarrowia lipolytica*, transcriptomics, lipid droplets, misfolded proteins, lipid classes, fatty acid toxicity, unfolded protein response, lipotoxicity

## Abstract

*Yarrowia lipolytica* naturally saves carbon excess as storage lipids. Engineering efforts allow redirecting the high precursor flux required for lipid synthesis towards added-value chemicals such as polyketides, flavonoids, and terpenoids. To redirect precursor flux from storage lipids to other products, four genes involved in triacylglycerol and sterol ester synthesis (*DGA1, DGA2, LRO1, ARE1*) can be deleted. To elucidate the effect of the deletions on cell physiology and regulation, we performed chemostat cultivations under carbon and nitrogen limitation, followed by transcriptome analysis. We found that storage lipid-free cells show an enrichment of the unfolded protein response, and several biological processes related to protein refolding and degradation are enriched. Additionally, storage lipid-free cells show an altered lipid class distribution with an abundance of potentially cytotoxic free fatty acids under nitrogen limitation. Our findings not only highlight the importance of lipid metabolism on cell physiology and proteostasis, but can also aid the development of improved chassy strains of *Y. lipolytica* for commodity chemical production.

**Highlights:**

- Physiological and transcriptomic characterization of storage lipid free (Q4) strain
- Storage lipid free strain shows an increased free fatty acid fraction on nitrogen limitation
- Storage lipid free strain is more sensitive towards fatty acid supplementation
- Unfolded protein response, chaperones, and ubiquitin are enriched in the storage lipid free strain

## Introduction

The initial interest in the oleaginous yeast *Yarrowia lipolytica* was focused on its ability to secrete lipases and its lipid production capacity^1^. *Y. lipolytica* can produce up to 80% of its dry weight as lipids thanks to metabolic engineering and media optimization^2^. The lipids produced by *Y. lipolytica* find applications as biofuels, nutritional supplements, cosmetic additives, and vegetable oil substitutes^3,4^. Lipid biosynthesis requires high amounts of acetyl-CoA. High amounts of acetyl-CoA can be redirected from lipid production to commodity and added-value chemicals production through metabolic engineering. Consequently, the interest in *Y. lipolytica* started to encompass the production of flavonoids (naringenin^5^, eriodictyol^6^, taxifolin^6^) polyketides (triacetic lactone^7^, resveratrol^5^), and terpenoids (lycopene^8^, β-Carotene^9^, limonene^10^).

Under nitrogen limitation (N-lim) *Y. lipolytica* has a high acetyl-CoA flux which is directed towards lipid accumulation^11,12^. Under N-lim the activity of the adenosine monophosphate deaminase (AMP deaminase) is increased, resulting in low AMP levels. Low AMP levels inhibit isocitrate dehydrogenase in the TCA cycle, resulting in citrate accumulation in the mitochondria. Mitochondrial citrate is shuttled into the cytosol by the malate/citrate transferase where it is cleaved into acetyl-CoA by the ATP-citrate lyase. Acetyl-CoA can enter lipid metabolism and can be incorporated into storage lipids. Storage lipids in *Y. lipolytica* are mainly formed by triacylglycerols (TAGs) and small amounts (<5%) of sterol esters (SE)^13^. Storage lipids are produced on the endoplasmic reticulum (ER) membrane and accumulate in lipid droplets (**Figure 1**)^14,15^.

**Figure 1:**
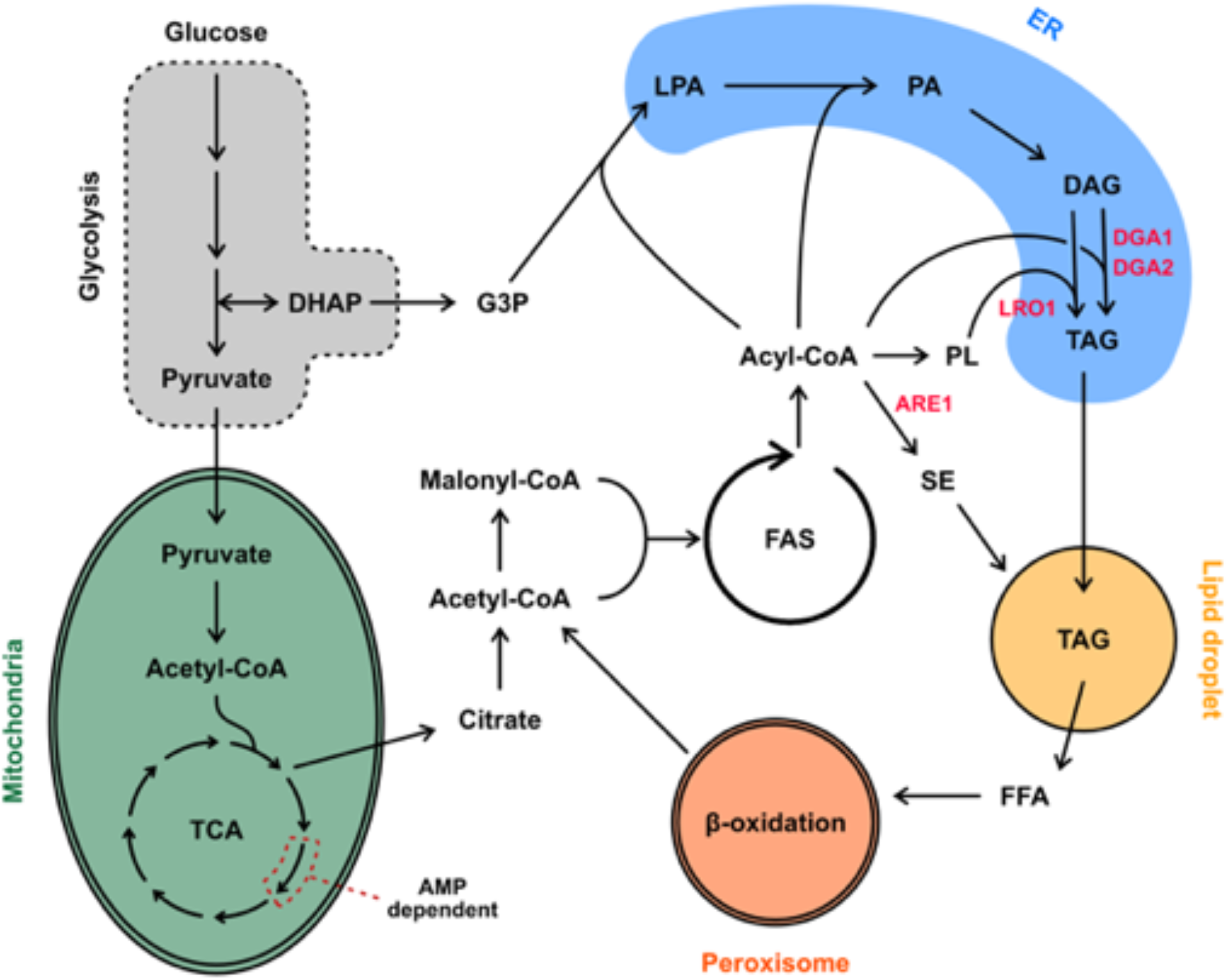
Overview of lipid metabolism in *Y. lipolytica*. In red are the genes that have been deleted in JFYL007 (also referred to as Q4 strain) to abolish storage lipid accumulation. Adapted from ^19^. DAG: diacylglycerol; DHAP: dihydroxyacetone phosphate; ER: endoplasmic reticulum; FAS: Fatty acid synthase; FFA: free fatty acid; G3P: Glyceraldehyde 3-phosphate; LPA: lysophosphatidic acid; PA: Phosphatidic acid; PL: phospholipid; SE: sterol ester; TAG: triacylglycerol; TCA: tricarboxylic acid cycle.

Storage lipid production can be prevented by deleting four genes, namely *DGA1* (YALI1_E38810g), *DGA2* (YALI1_D10264g), *LRO1* (YALI1_E20049g), and *ARE1* (YALI1_F09747g)^13^. *DGA1* and *DGA2* catalyse the final step of TAG formation and use acyl-CoA as a substrate to convert diacylglycerols (DGAs) into TAGs^13,16^. *DGA2* has also been reported to affect the size and morphology of lipid droplets^13^. *LRO1* codes for a triacylglycerol synthase that is acyl-CoA independent and uses phospholipids as acyl-donors to convert DAGs into TAGs^16^. The Are1p of *Y. lipolytica* is essential for sterol esterification, as deletion of the encoding gene (*ARE1*) completely abolished SE synthesis^13^. Decreasing or inhibiting storage lipid accumulation has been shown to increase production of added-value and commodity chemicals. Shi et al.^17^ showed that deleting *DGA1* and *DGA2* increases β-farnesene titres by 56%.

A *Y. lipolytica* strain carrying *DGA1, DGA2, LRO1*, and *ARE1* deletions (referred to as Q4 strain) lacks storage lipid accumulation^13,18^. While abolishing storage lipid accumulation is a justifiable strategy to increase the production of desired chemicals in *Y. lipolytica*, the consequences on cell physiology and regulation have not been investigated yet. To elucidate the effect of abolishing storage lipid synthesis, we performed chemostat cultivations on the wild-type strain and the Q4 strain, under both carbon limitation (C-lim, C/N ratio 3) and nitrogen limitation (N-lim, C/N ratio 116). We monitored physiological parameters, the abundance, composition, and class distribution of lipids, and we performed transcriptome analysis to assess how the deletions affect cell regulation and homeostasis.

## Results

### Gene deletions affect cell physiology and lipid chain composition in nitrogen limitation

Nutrient limitation has a major impact on gene expression in *Y. lipolytica*^20^, and under N-lim high flux through acetyl-CoA enables lipid accumulation^11,12^. To select suitable C/N ratios for chemostat fermentations, we performed shake-flask cultivations with different C/N ratios. We grew *Y. lipolytica* in media with varying glucose concentrations while maintaining the same nitrogen concentration to test C/N ratios between 1.45 and 20 and measured the OD^600^ after 72 hours. Based on the results (**Figure S1**) we selected C/N ratio of 3 (C-lim, lipid accumulation not stimulated), and a C/N ratio of 116 (N-lim, lipid accumulation stimulated).

To assess if cell physiology changes are related to (i) gene deletions; (ii) the C/N ratio; or (iii) a combination of both factors, we cultivated *Y. lipolytica* in chemostats under N-lim and C-lim. The pH-controlled chemostat fermentations were performed on OKYL029, displaying wild-type lipid phenotype, and on JFYL007, referred to as Q4, carrying four lipid-genes deletions (*∆dga1, ∆dga2, ∆lro1, ∆are1*), and previously used to study lipid metabolism and the role of acyltransferases^13,21,22^. In both strains hyphae formation was abolished by deleting the gene *MHY1*^23^. We picked pH-controlled chemostat cultivations to ensure a highly controlled environment that increases reproducibility. Additionally, it is possible to control the growth rate by varying the dilution rate, reducing growth-related variability, and ensuring comparable results between strains with different growth dynamics (**Figure S2**). After at least five-volume changes in chemostat fermentation, we measured the cell physiology parameters such as biomass, lipid content and composition, biomass and lipid yield, and glucose consumption.

In C-lim cell dry weight (CDW) and lipid content were unaffected by the four deletions in the Q4 strain (**Figure 2**). In C-lim the two strains showed similar biomass yields, lipid yields, and specific glucose uptake rates (r-glucose) (**Figure 2**).

**Figure 2:**
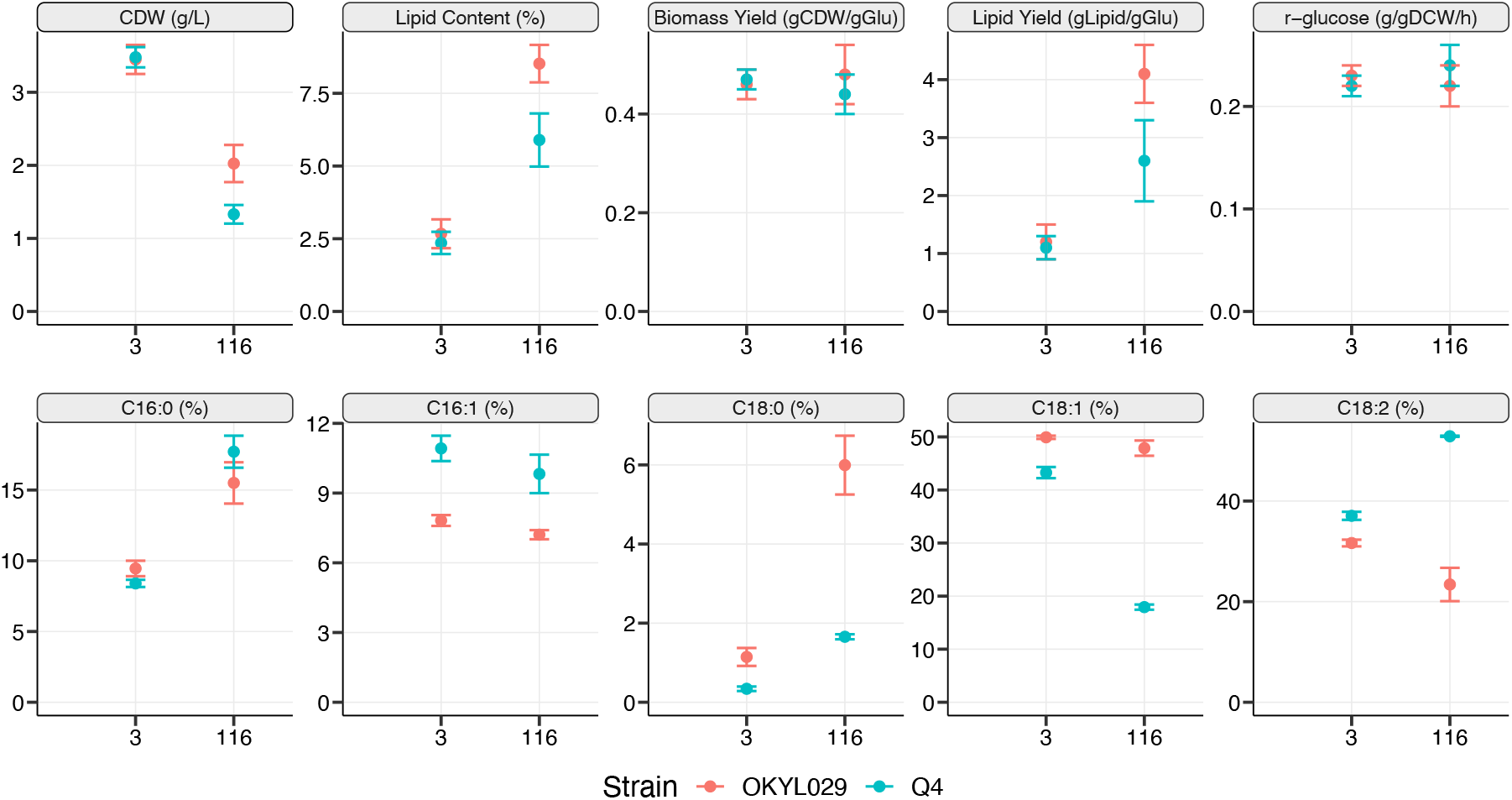
Physiological and lipid composition changes of the strains OKYL029 and Q4 in C-lim (C/N ratio 3) and N-lim (C/N ratio 116). Lipid content is calculated as % of the lipids on the CDW, and the strains’ fatty acid composition is calculated as the % of each chain length on the total amount of lipids. Displayed are the average (dot) and standard deviation (error bar) of at least three replicates.

Even though the Q4 strain is carrying four deletions in genes involved in TAG and SE synthesis, the lipid content in the Q4 strain increased when cultivated under N-lim compared to C-lim (5.9% on N-lim and 2.3% C-lim). Since TAG and SE synthesis are knocked out, the Q4 strain likely accumulates DAGs.

Under N-lim both the CDW and the lipid content were negatively affected by the deletions. The CDW and the lipid content decreased 33% and 32% in the Q4 strain compared to OKYL029, respectively. Cells grown under N-lim were dyed with Bodipy® Lipid Probe and lipid droplets (LD) were visible in the wild-type strain, while the Q4 strain showed no visible LDs (**Figure S3**). The lipid yield in Q4 was 37% lower than in the OKYL029 strain. Both strains showed similar biomass yields (0.44 gDCW/gGlu in Q4 and 0.48 gDCW/gGLU in OKYL029) and the same specific glucose uptake rate (r^glucose^, 0.23 g/gDCWh in Q4 and 0.22 g/gDCWh in OKYL029).

Since four genes involved in lipid synthesis are deleted in the Q4 strain, we investigated their effect on the abundance and chain length of the lipids synthesized. For that, we converted the lipids to fatty acid methyl ester (FAME) and analysed them using Gas Chromatography Mass Spectrometry (GC-MS). We found that the quadruple deletions affected the lipid composition in both C/N ratios. Regardless of the C/N ratio the difference between strains in the C16:0 fraction is not statistically significant (p-value > 0.01). All the other fatty acids (FAs) (C16:1, C18:0, C18:1, C18:2) are significantly different (p-value < 0.01) between the two strains, in both C/N ratios. The entity of the change is generally bigger in N-lim, but the direction of the change is the same (either more abundant in both C/N ratios or less abundant in both C/N ratios).

Overall, these results indicate that the gene deletions have a major effect on cell physiology and lipid chain composition under nitrogen limitation. On the other hand, under C-lim cell physiology resulted mainly unchanged and we could only detect small changes in the FA chain length of the lipids. Since the Q4 strain showed altered lipid chain abundances compared to the wild-type strain, we performed a more detailed lipid analysis (solid-phase extraction). We separated the different lipid classes to validate if differences are in membrane lipids, storage lipids, or free fatty acids.

### Q4 strain shows higher free fatty acid and phospholipid fractions on nitrogen limitation

The FAME analysis only provides insight into the changes in the total lipid composition of the cell but does not discriminate between different lipid classes, such as neutral lipids (NL, consisting of DAGs, TAGs, and SE), free fatty acids (FFA), and phospholipids (PL, mainly found in membranes). Therefore, we performed a solid-phase extraction (SPE)^24^ which separates the NL, FFA, and PL fractions and analyzes the fatty acid distribution of each fraction.

Under C-lim there are no statistically significant differences (p-value > 0.01) between the Q4 and the wild-type strain. Regardless of the strain, 73-77% of the lipids are PL (**Figure 3)**, followed by 16-23% of NL and 4-7% of FFA. The chain length distribution of the PL fraction shows statistically significant differences (p-value < 0.01), but the entity of the change is less than 5% (**Figure 3**). We detected no significant differences in the chain length distribution of NL (**Figure 3B**). In the FFA fraction the Q4 strain has higher percentages of unsaturated lipids (C16:1, C18:1, and C18:2), while the saturated lipids (C16:0 and C18:0) are more abundant in the wild-type strain OKYL029 (**Figure 3**). However, it is worth noting that the FFA fraction only contributes minorly to the overall lipid content (7% in the Q4 strain and 4% in OKYL029 strain).

**Figure 3:**
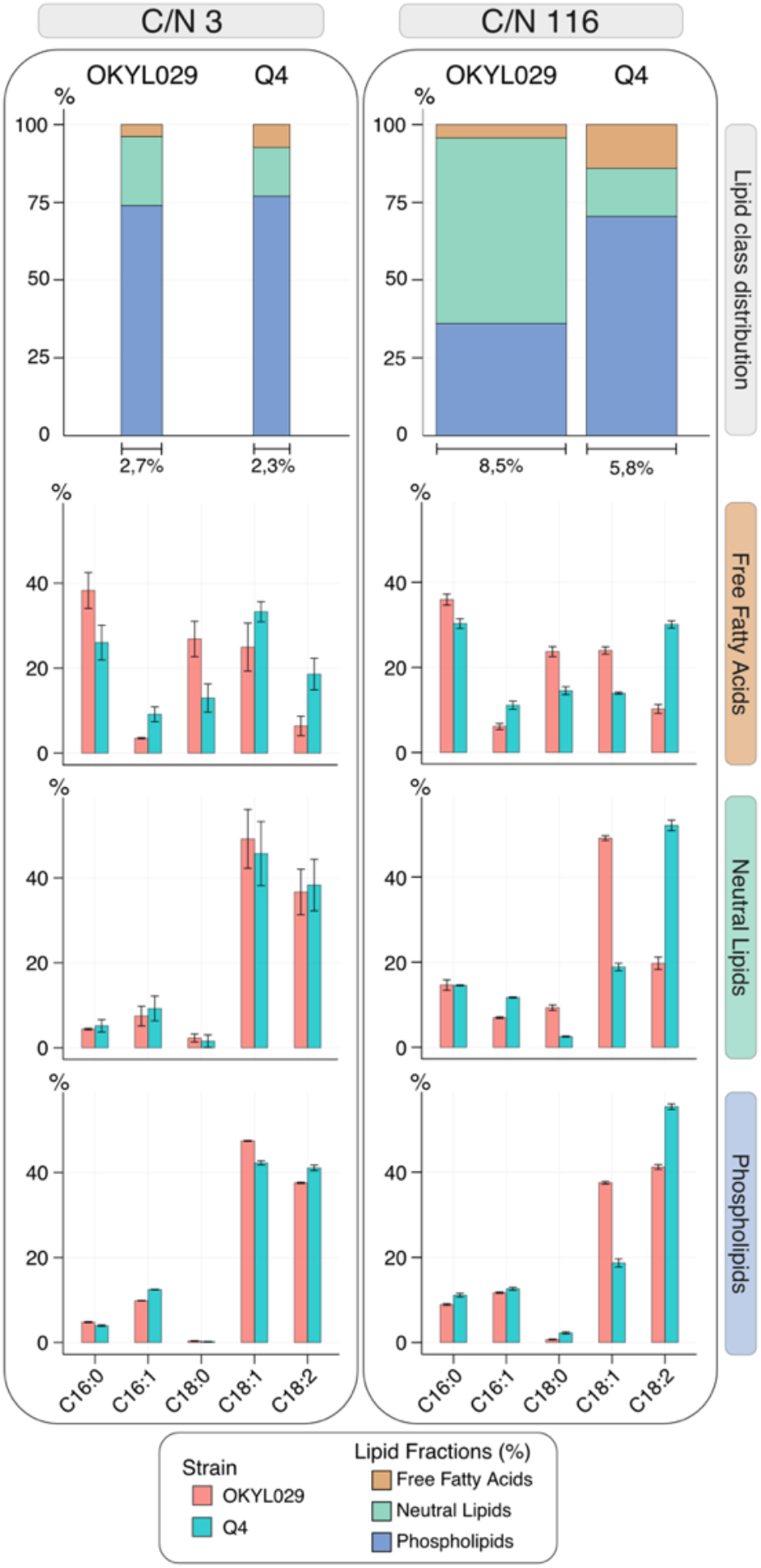
Solid phase extraction (SPE) of Q4 and OKYL029 in C-lim and N-lim conditions. Stacked bar charts (lipid class distribution) represent the share of each lipid class detected by SPE over the total amount of lipids present in the cell. The strains OKYL029 and Q4 were analyzed in C-lim and N-lim. The bar chart area is proportional to the total lipid content of the cell. The bottom three bar charts represent the fatty acid composition of each lipid fraction (free fatty acids, neutral lipids, and phospholipids), calculated as the % of each chain length on the amount of lipids in that specific lipid class. Displayed is the average and standard deviation of at least three replicates.

In the OKYL029 strain the NL fraction accounts for only 23% of the total cell lipids in C-lim, compared to 60% in N-lim (**Figure 3**), while the PL fraction is increased (from 36% to 73%). The FFA fraction only accounts for 4% of the total cell lipids in both C/N ratios. In the Q4 strain the share of the NL fraction remained similar (15%-16%) in both C-lim and N-lim (**Figure 3**). The PL fraction decreased 7% (77% in C-lim, 70% in N-lim), and the FFA fraction doubled (7% in C-lim to 15% in N-lim). In N-lim, regardless of the lipid fraction, C16:1 and C18:2 have a higher share in the Q4 strain than the wild-type strain. The C18:1 is more abundant in the wild type, regardless of the lipid class. The saturated fatty acids (C16:0 and C18:0) only show minor changes between strains.

These observations indicate that under C-lim the Q4 strain only shows minor differences in the lipid fractions compared to the wild-type strain, confirming the results from our previous FAME analysis. On the other hand, under N-lim, the Q4 strain shows higher free fatty acid and phospholipid fractions. These results indicate that the deletions in TAG and SE synthesis mainly affect the cell phenotype when lipid synthesis is stimulated under N-lim.

### Fatty acid supplementation affects the growth of the Q4 strain

Previous studies in *S. cerevisiae* suggested a crucial interplay between lipid synthesis, lipid droplets, and protein homeostasis^25^. It has been reported that a storage lipid-free Q4 strain of *S. cerevisiae* (∆*ARE1*, ∆*ARE2*, ∆*DGA1*, ∆*LRO1*) shows higher sensibility towards unsaturated fatty acid supplementation, indicating the important role TAGs have in fatty acid buffering and detoxification^26^.

To assess the sensitivity of *Y. lipolytica* towards unsaturated fatty acids, we calculated the molar concentrations of each fatty acid (C16:0, C16:1, C18:0, C18:1, C18:2) both in the FFA fraction and in the total lipids (**Figure S4**). The highest fatty acid concentration calculated in the FFA fraction is 7 μm (C16:0). When considering the total lipids, the highest concentration measured is 1.5 mM (C18:1). We tested these concentrations but no effect was visible (data not shown). Therefore we decided to test the highest concentrations that solubility allowed in our experimental setup, and supplemented cultivations with up to 8 mM of unsaturated fatty acids and up to 1 mM of saturated fatty acids (**Figure 4**).

**Figure 4:**
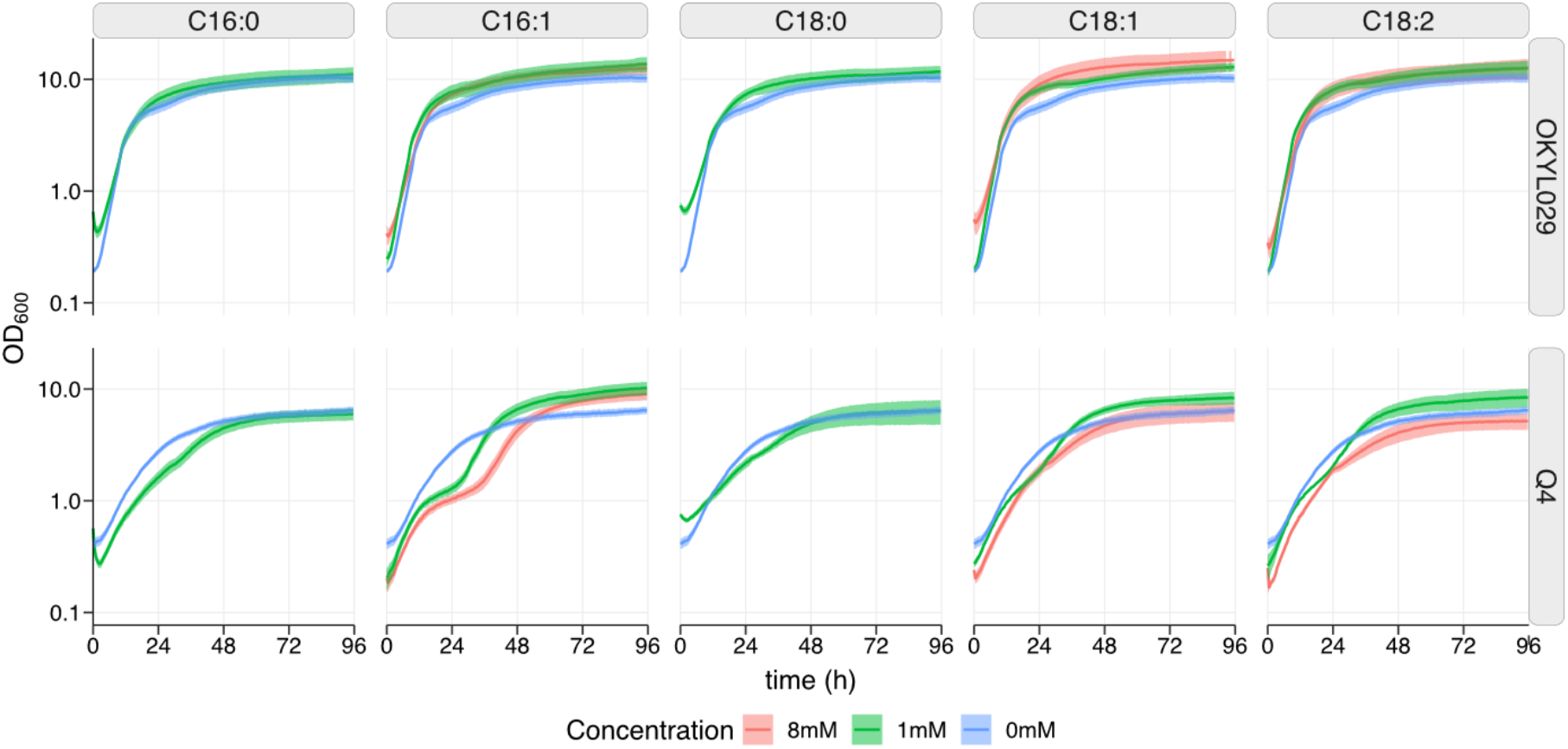
Growth curves of *Y. lipolytica* strains Q4 and OKYL029 on delft media containing 2% ethanol and 1% tween-20. The media was supplemented with different concentrations of fatty acid. Strains were cultured in 96-well plates and the OD_600_ was measured with the growth profiler every 30 minutes. The lines and shadows represent the average and standard deviation of five replicates.

Overall, the Q4 strain is more sensitive to fatty acids, while the wild-type strain was unaffected even by high concentrations. For example, supplementation of the unsaturated palmitoleic acid (C16:1) did not affect OKYL029, while it induced a different growth profile in the Q4 strain. The Q4 strain exhibited two exponential growth phases that were interrupted by a period of reduced growth. Even though FFA affects the growth of the Q4 strain, the strain is less sensitive to fatty acid supplementation compared to *S. cerevisiae*, in which concentrations above 0.5 mM delay or inhibit growth^27^.

### Gene deletions affect gene expression in nitrogen limitation but not in carbon limitation

To elucidate how the four deletions impact cell regulation, we performed transcriptomic analysis (RNA-seq) on pH-controlled chemostat samples from the Q4 and OKYL029 strains, in carbon and nitrogen limitation.

Similarities and dissimilarities between samples were assessed with principal component analysis (PCA) (**Figure 5A)**. The first two principal components (PC) respectively account for 55% and 15% of the total variance of the RNA-seq dataset. The PCA shows that the two main experimental factors, C/N ratio and strain background, separate the samples and account for most of the variability. Additionally, samples in C-lim cluster together, regardless of their genetic background, while samples are separated by genetic background in N-lim, when lipid accumulation is stimulated. These results align well both with phenotype and lipid measurements, where in C-lim both strains are very similar, while in N-lim the strains show major differences.

**Figure 5:**
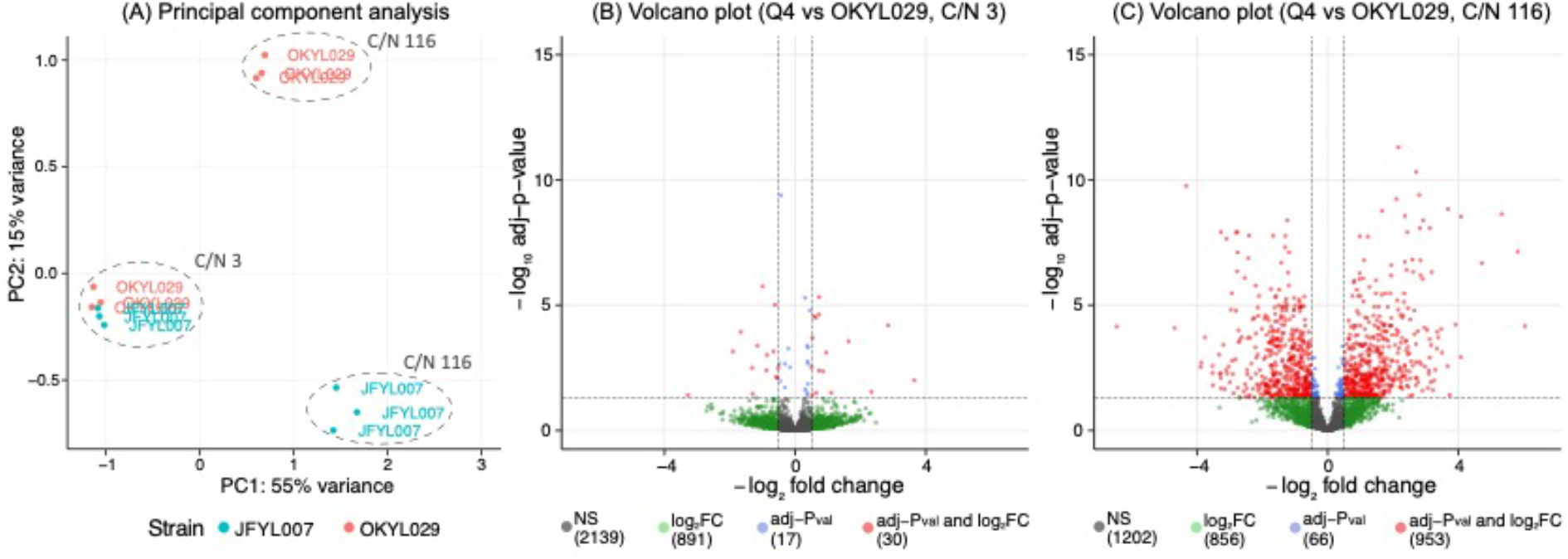
RNA-sequencing of *Y. lipolytica* strains Q4 and OKYL029 in carbon (C/N ratio 3) and nitrogen (C/N ratio 116) limitation. Panel A: principal component analysis. Volcano plots for samples in carbon (B) and nitrogen limitation (C). NS: non-significative genes. Log_2_FC: genes with an absolute fold change greater than 0.5. Adj-p-value: genes with an adjusted-p-value below 0.05. Log_2_FC and adj-p-val: genes with both adjusted-p-value below 0.05 and absolute log_2_FC greater than 0.5.

To identify changes in expression levels and to identify which genes are differentially expressed between samples, we performed differential gene expression analysis and compared the Q4 strain with the OKYL029 in C-lim and N-lim. As expected, after p-value adjustment, for samples clustering together in C-lim we only detected 30 differentially expressed genes, either upregulated or downregulated (absolute log_2_ fold change greater than 0.5 and adjusted p-value below 0.05) (**Figure 5B**). For each differentially expressed gene we checked the annotated function on UniProt. Unfortunately, most genes have an unknown function and we only found 6 genes associated with a protein function (**Table S1**). Three of these genes are related to lipid metabolism (“triacylglycerol lipase”, “glycerophosphocholine phosphodiesterase”, “Glycolipid 2-alpha-mannosyltransferase-domain-containing protein”) and might contribute to the small differences we observed in lipid composition between strains in carbon limiting conditions.

The major difference between strains is in N-lim when nitrogen depletion triggers lipid accumulation. As expected from the PCA results, we see major differences between strains. By comparing the Q4 strain to OKYL029 we found a total of 953 differentially expressed genes (absolute log_2_ fold change above 0.5 and adjusted p-value below 0.05) (**Figure 5C**). Out of the total 953 differentially expressed genes, 390 have a function annotated on UniProt (**Table S2**).

In summary, the low variance identified in the PCA and the low number of genes differentially expressed indicate that the deletions have minimal effect on the overall transcriptome in C-lim. On the other hand, the variance identified in the PCA and the high number of differentially expressed genes indicates a major effect of the deletions in N-lim.

### Gene set analysis reveals upregulation of misfolded protein gene sets

The high number of differentially expressed genes in the Q4 strain under N-lim hinders single-gene analysis. To draw biological conclusions, we performed a gene set analysis (GSA). In a GSA the gene sets are defined based on prior biological knowledge to determine whether a defined gene set shows statistically significant differences between samples^28^. Gene sets can be defined using gene ontology (GO) terms^29^. GO terms are generally divided into biological process, molecular function, and cellular component. For each of these levels, we performed a gene set analysis with the R package PIANO^30^, using gene-level statistics calculated with TMM-normalized gene counts (**Figure 6**).

**Figure 6:**
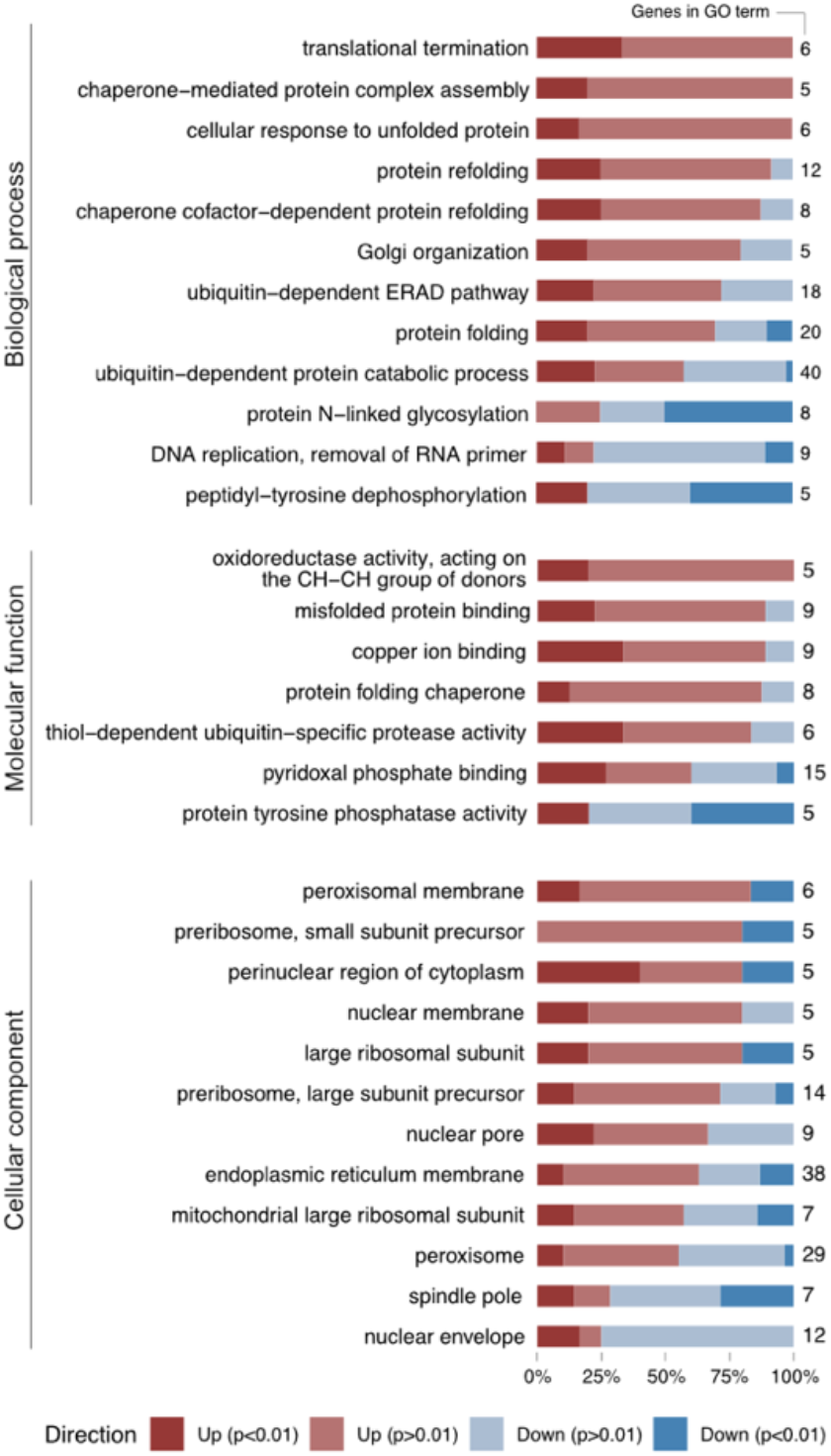
Gene set analysis (GSA) of Q4 vs OKYL029 in N-lim (C/N ratio 116). Gene sets are defined by GO terms (biological process, molecular function, cellular component). For each gene set that is significantly enriched, the direction of the relative changes in RNA levels (positive or negative fold change) is shown, and the genes in the gene sets are marked based on significative or non-significative adjusted p-value (cut-off 0.05). Genes are considered up or down in the Q4 strain, and the OKYL029 strain is the reference strain. The total number of genes in each gene set is reported on the right.

A biological process represents a specific objective that the organism is genetically programmed to achieve, for example, the biological process of cell division, and is carried out by specific gene products in a regulated manner^31^. In the biological process, we found 12 gene ontology (GO) terms that are significantly enriched in the Q4 strain. Among them, broad GO terms such as “protein folding”, “protein refolding”, and “cellular response to unfolded protein” are enriched. Two GO terms related to chaperones and chaperones activity are enriched in the Q4 strain, namely “chaperone-mediated protein complex assembly” and “chaperone cofactor-dependent protein refolding”. Chaperones are proteins that assist conformational folding of proteins during or after synthesis, and after partial denaturation^32^. An enrichment of chaperone-related processes indicates that the cell has activated mechanisms to handle folding stress. We also found that ubiquitin-related GO terms are enriched. The “ubiquitin dependent ERAD pathway” targets ER-resident proteins for degradation to the cytoplasmic proteasome^33^. The “ubiquitin-dependent protein catabolic process” is a group of reactions resulting in protein degradation, and is initiated by multiple ubiquitin groups binding to the protein targeted for degradation^34^. Two other GO terms (“Golgi organization” and “protein N-linked glycosylation”) are enriched in the Q4 strain and can be linked to unfolded protein response. Most genes in the GO term “Golgi organization” are upregulated. Signalling to transcribe genes for Golgi organization is active, suggesting that a proper Golgi organization is missing. The genes in the “protein N-linked glycosylation” are mainly downregulated, suggesting that transcription of genes responsible for glycosylation is not required. This can be due to newly synthesized proteins being misfolded and targeted for degradation instead of being transported to the Golgi apparatus for glycosylation. Two GO terms (“translation termination” and “DNA replication removal of RNA primer”) are related to cell growth and replication. For the GO term “peptidyl-tyrosine dephosphorylation” the annotation is insufficient to reconstruct its precise biological significance, but dephosphorylation and phosphorylation of tyrosine residues modulate the enzymatic activity and creates binding sites for the recruitment of downstream signalling proteins^35^.

A molecular function term describes activities that occur at the molecular levels and are carried out by individual gene products or by molecular complexes composed of multiple gene products^31^. In the molecular function, we found seven significantly enriched GO terms in the Q4 strain. Three GO terms (“misfolded protein binding”, “protein folding chaperone”, “thiol-dependent ubiquitin-specific protease activity”) are broadly related to misfolded protein response. Two GO terms (“copper ion binding” and “oxidoreductase activity, acting on the CH-CH group of donors”) play important roles in many biochemical processes such as oxidation, dioxygen transport and electron transfers. The lack of further annotation prevents the reconstruction of more detailed molecular functions for these gene sets. “Pyridoxal phosphate binding” is related to pyridoxal phosphate, the active form of the B6 vitamin, that acts as a cofactor for enzymes involved in the biosynthesis of amino acids and amino acid-derived metabolites^36^. “Protein tyrosine phosphorylation” is a post-translational modification that creates novel recognition motifs for protein interactions and cellular localization^35^.

Cellular component is the location occupied by a macromolecular machine when it carries out a molecular function^31^. It can be relative to cellular compartments, structures, or macromolecular complexes (e.g., the ribosome). We found 12 differentially enriched GO terms in the Q4 strain, that are mainly related to transcription and translation. Four GO terms are related to the nucleus (“perinuclear region of cytoplasm”, “nuclear membrane”, “nuclear pore”, “nuclear envelope”), three are related to ribosomes (“preribosome, small subunit precursor”, “large ribosomal subunit”, “preribosome, large subunit precursor”), and two are related to the peroxisomes (“peroxisomal membrane”, “peroxisome”). Additionally, we saw an enrichment of the GO terms “endoplasmic reticulum membrane”, “mitochondrial large ribosomal subunit”, and “spindle pole”.

Our results show that many GO terms related to unfolded protein response (UPR), chaperones, and ubiquitin are enriched in the Q4 strain. Molecular functions and cellular components related to UPR response are also affected, indicating that the cell is experiencing folding stress, and protein synthesis and functionality are affected by the four deletions. These results suggest a link between physiological lipid production and protein homeostasis.

## Discussion

Deleting lipid genes has proven a valid strategy to boost added value and commodity chemicals production in *Y. lipolytica*^17^, but it would be undesired if those deletions would negatively impact cell physiology and reduce cell resilience and robustness. In this context, our goal was to elucidate the consequences of deleting four genes involved in lipid metabolism (*DGA1, DGA2, ARE1, LRO1*). We cultivated the Q4 strain (JFYL007) carrying the four deletions and the wild-type strain (OKYL029) either in carbon or nitrogen-limiting conditions and performed RNA-sequencing. We showed that in C-lim physiological parameters and the transcriptome only show a few minor changes, while in N-lim, when the carbon flux towards lipid synthesis is high, cell physiology and the transcriptome are strongly affected by the deletions. Furthermore, we show that in N-lim the lipid organization of the Q4 strain is altered, and multiple GO terms related to unfolded protein response (UPR) are enriched in the transcriptome.

In N-lim the SPE analysis revealed that the lipid distribution in neutral lipids (NL), phospholipids (PL), and free fatty acids (FFA) are affected in the Q4 strain. While the wild-type strain showed a predominant NL fraction, the Q4 strain has reduced amounts of NL, and PL are predominant. The FFA fraction in the Q4 strain is three times larger than in the wild-type strain, indicating that the Q4 strain is synthesising FFA but lacks the ability to incorporate them in TAGs. Lipid homeostasis is maintained by balancing NL synthesis and lipid turnover^37^. To avoid possible toxic and membrane-disturbing effects, FFA are stored as NL, which are biologically inert^37^. In *Y. lipolytica* the NL fraction mainly contains TAGs, and only small amounts of SE^13^. The Q4 strain lacks four genes responsible for TAG and SE synthesis. In C-lim lipid synthesis is not stimulated and the genotypical difference between the Q4 strain and the wild-type strain is not visible in the phenotype. In N-lim the flux through the four enzymes of the lipid accumulation pathway is high^11,12^ and their absence prevents FFA to be incorporated into TAGs.

A storage lipid-free Q4 strain of *S. cerevisiae* (∆*ARE1*, ∆*ARE2*, ∆*DGA1*, ∆*LRO1*) shows high sensibility towards FFA supplementation, suggesting the important role TAGs play in fatty acid buffering and detoxification^26^. FFA could act as detergents, interfering with membrane integrity, or could be incorporated into lipid species that are cytotoxic at high levels, such as ceramide, acylcarnitine and diacylglycerol^25^. In wild-type strains, to prevent lipotoxicity, excess FFA are incorporated into TAGs, which are in turn stored into LDs^2,37^. The Q4 strain of *Y. lipolytica* cannot synthesize LDs^22^ (**Figure S3**). The higher levels of FFA detected in the Q4 strain under N-lim, and their reported cytotoxicity may be the reason for the lower CDW reached by the Q4 strain during the fermentation, which was 33% lower compared to the wild-type strain. We investigated *Y. lipolytica’s* sensitivity towards fatty acids (**Figure 4**) and showed that the Q4 strain is more sensitive to high concentrations of unsaturated fatty acids than the wild-type strain. The growth of the Q4 strain was affected by FFA, but the strain was able to grow in media supplemented with 8 mM FFA. The Q4 strain of *Y. lipolytica* is less sentive to fatty acid supplementation then the Q4 strain of *S. cerevisiae*, in which concentrations of 0.5 mM delay or inhibit growth^27^. Additionally, if a downstream pathway is integrated in the Q4 strain the acyl-CoA flux would be redirected to other products, which should prevent accumulation of FFAs.

RNA-sequencing and gene set analysis revealed an enrichment of multiple GO terms connected to the unfolded protein response (UPR) when the Q4 strain was cultivated under N-lim conditions. We found broad GO terms contributing to the UPR response and four GO terms related to chaperones and ubiquitin-dependent activities enriched in the Q4 strain. Chaperones are proteins that assist the conformational folding of proteins during or after synthesis, and after partial denaturation^32^. Ubiquitin-dependent activities are responsible for targeting proteins for degradation^33,34^. The Q4 strain shows enrichment of chaperone and ubiquitin-related related GO terms, indicating that the cells are experiencing folding stress. The enrichment of Golgi-related GO terms further supports this observation. Proteins are glycosylated in the Golgi apparatus before being targeted for delivery to their final destination^38^. The genes in the “protein N-linked glycosylation” GO term are mainly downregulated, suggesting that newly synthesized proteins might be misfolded and targeted for degradation before being transported to the Golgi apparatus for glycosylation. The genes of the “Golgi organization” GO term are mainly upregulated, suggesting that a proper Golgi organization might be lacking. Taken together, these results suggest that a major alteration in lipid metabolism affects protein synthesis and functionality.

The Q4 strain lacks the ability to synthesize LDs^22^ (**Figure S3**) and displays alterations in the quantity and lipid distribution, activation of the UPR response, and enrichment of several GO terms related to proteostasis. These observations indicate that cell homeostasis is linked with lipid droplet biology and functionality, as was previously shown in *S. cerevisiae*^39^. Lipids are synthesized and aggregate on the ER membrane, where they form lipid droplets (LD) that can bud from the ER membrane^40^. LDs have a neutral lipid core surrounded by a monolayer of phospholipids, usually associated with proteins^40^. LDs have been shown not only to act as lipid storage but also to prevent lipotoxicity by buffering fatty acid stress^26,41^ and to have an active role in membrane and organelle homeostasis^26,39,42^. LDs are important in starvation-induced autophagy^42,43^, clearance of inclusion bodies^44^, and, ultimately, in proteostasis^42,44^. Starvation-induced autophagy, for example during nitrogen starvation, is a physiological process involved in protein and organelle degradation in the lysosomes^43^; inclusion bodies are non-toxic and non-soluble aggregates of misfolded proteins. LDs participate in starvation-induced autophagy and physically associate with inclusion bodies, contributing to their degradation^44^. Proteostasis is the group of processes that regulate protein synthesis and degradation within the cell. LDs are involved in proteostasis mechanisms and have an important role in contributing to stress resistance, cell survival, and, ultimately, cell homeostasis. Deleting gene involved in lipid metabolism in *Y. lipolytica* results in cells that lack LDs, which are important organelles in cell homeostasis. This results in cells with altered lipid composition and proteome, and with upregulation of the UPR response.

To summarise, our work reveals the connection between lipid metabolism, protein regulation and folding, and cell homeostasis. Deleting the genes *DGA1, DGA2, LRO1*, and *ARE1* blocks TAG and SE accumulation in the Q4 strain, and results in the absence of LDs. LDs play an important role in lipid homeostasis and participate in the clearance of misfolded proteins and inclusion bodies. Lacking LDs, the Q4 strain not only shows altered lipid class distribution, but also an abundance of potentially cytotoxic FFA, and an enrichment of GO terms related to the UPR response, protein degradation, and Golgi activities. Our findings highlight the importance of lipid metabolism in *Y. lipolytica* and the implications of abolishing TAG synthesis on cell physiology and proteostasis. Our findings can aid the development of improved chassy strains of *Y. lipolytica* for commodity chemical production, where lipid synthesis is not abolished but downregulated to maintain cell robustness and physiological activities.

## Materials and methods

### Yeast strains

The strains used in the study are derived from the *Yarrowia lipolytica* strain ST6512^45^, which was derived from the W29 background strain (Y-63746 from the ARS Culture Collection, Peoria, USA (also known as ATCC20460/CBS7504). ST6512 has been engineered to harbor a KU70::Cas9-DsdA system, which allows for marker-free genomic engineering using the EasyCloneYALI toolbox^46^. The strain OKYL029 (ST6512 ∆mhy1) has a deletion of the MHY1 gene to prevent stress-induced hyphae formation^23^. The strain OKYL049 (ST6512 + E1::pTef1in + DGA1 + tPEX20 Δare1 Δmhy1) is an obese strain that carries overexpression of the DGA1 gene and a deletion of the ARE1 gene to increase TAG accumulation and abolish sterol ester formation^47^. To prevent hyphae formation, the MHY1 gene is also deleted in this strain. The Q4 strain JFYL007 (ST6512 ∆mhy1 ∆are1 ∆lro1 ∆dga1 ∆dga2) is a low-lipid accumulating strain that carries deletions of the ARE1, LRO1, DGA1, and DGA2 genes to decrease TAG accumulation, and a deletion of the MHY1 gene to prevent hyphae formation^18^. The gene annotation for these strains follows the YALI1 system, but translation into YALI0/CLIB122 can be done with the **S2** table of ^48^.

### Shake flask cultivation

To select suitable C/N ratios for chemostat cultivations, *Y. lipolytica* was grown for 5 days in delft media containing 7.5 g/L ammonium sulphate [Sigma Aldrich, 7783-20-2], 14.4 g/L magnesium sulphate heptahydrate [Merck, 10034-99-8], 0.5 g/L potassium dihydrogen phosphate [Sigma Aldrich, 7778-77-0], 1 mL of trace metals solution, and 1 ml of vitamin solution. The pH of the media was set at 5.0 by potassium hydroxide [Avantor, VWR, 1310-58-3] addition. The glucose [Avantor, VWR, 14431-43-7] concentration was varied between 3 g/L and 60 g/L to obtain C/N ratios between 1 and 20.

Trace metal solution consisted of 3 g/L Iron(II) sulfate heptahydrate (FeSO_4_*7 H_2_O) [Sigma Aldrich, 7782-63-0], 4.5 g/L Zinc sulfate heptahydrate (ZnSO_4_*7 H_2_O) [Sigma Aldrich, 7446-20-0], 4.5 g/L Calcium chloride dihydrate (CaCl_2_*2 H_2_O) [Sigma Aldrich, 10035-04-8], 1.0 g/L Manganese(II) chloride tetrahydrate (MnCl_2_*4 H_2_O) [Sigma Aldrich, 13446-34-9], 300 mg/L Cobalt(II) chloride hexahydrate (CoCl_2_*6 H_2_O) [Sigma Aldrich, 7791-13-1], 300 mg/L Copper(II) sulfate pentahydrate (CuSO_4_*5 H_2_O) [Sigma Aldrich, 7758-99-8], 400 mg/L Sodium molybdate dihydrate (Na_2_MoO4*2 H_2_O) [Sigma Aldrich, 10102-40-6, 1.0 g/L Boric acid (H_3_BO_3_) [Sigma Aldrich, 10043-35-3], 100 mg/L Potassium iodide (KI) [Sigma Aldrich, 7681-11-0], and 19 g/L Disodium ethylenediaminetetraacetate dihydrate (Na_2_EDTA*2 H_2_O) [Sigma Aldrich, 6381-92-6]. Vitamin solution consisted of 50 mg/L d-biotin [Sigma Aldrich, 58-85-5], 1.0 g/L D-pantothenic acid hemicalcium salt [Sigma Aldrich, 137-08-6], 1.0 g/L thiamin-HCl [Sigma Aldrich, 67-03-8], 1.0 g/L pyridoxin-HCl [Sigma Aldrich, 58-56-0], 1.0 g/L nicotinic acid [Sigma Aldrich, 59-67-6], 0.2 g/L 4-aminobenzoic acid [Sigma Aldrich, 150-13-0], 25 g/L myo-inositol [Sigma Aldrich, 87-89-8].

### Growth profiler

To investigate fatty acid toxicity, *Y. lipolytica* strains OKYL029, and JFYL007 (Q4) were cultivated in 96-well plates, at 30 °C and 200 rpm. Growth performances were determined with Growth Profiler 960 (Enzyscreen B.V., Heemstede, The Netherlands) using a standard sandwich cover with pins (CR1396b) with OD_600_ measurement every 30 min. To determine the growth performances, cells were grown in 150 μl of delft media (N-lim, C/N 116) containing 1% Tween-20, 2% ethanol, and fatty acids (C16:0, C16:1, C18:0, C18:1, C18:2) at a final concentration of 8 mM or 1 mM. The controls were grown in 150 μl of delft media (N-lim, C/N 116) containing 1% tween-20 and 2% ethanol. The starting OD_600_ for cell cultivation was 0.1. Each experiment was carried out in quintuplicate. Growth curves represent the average of quintuplicates, and error bars are represented as shadowed areas.

### Bioreactor and chemostat cultivation

Chemostat cultivations were performed as described in ^49^. Cultivations were carried out in DasGip 1-L stirrer-pro vessels (Eppendorf, Jülich, Germany). The working volume was 500 mL, the temperature was kept at 28 °C, and the agitation was set at 600 rpm. To ensure aerobic conditions, sterile airflow was set at 1 vvm (= 30 Lh^-1^) and dissolved oxygen was monitored with DO probes (Mettler Toledo, Switzerland). The pH was monitored with a pH sensor (Mettler Toledo, Switzerland) and maintained at 5.0 ± 0.05 by automatic addition of 2M KOH. Batch cultivations were performed with the same media as chemostat cultivations and cell growth was monitored via the CO_2_ exhaust gas. After the cells left the exponential growth phase, the constant feed was initiated to obtain steady-state cultivation with a dilution rate of 0.10 h^−1^. The working volume of 500 mL was maintained using an overflow pump. Samples for transcriptome analysis were taken after at least 4 residence times of steady-state growth, and each condition was cultivated at least in triplicates.

Chemostat cultivations with C/N ratio 3 (C-lim) were performed in delft media containing 5.28 g/L of ammonium sulphate, 7.92 g/L glucose, 0.5 g/L magnesium sulphate heptahydrate, 3 g/L monopotassium phosphate, 1 mL of trace metals solution, and 1 mL of vitamin solution. Trace metal and vitamin solutions have the same composition as in shake flask cultivations. The pH was set at 5 with 2M KOH and was kept at 5 during the fermentation by automatic KOH addition.

Chemostat cultivations with C/N ratio 116 (N-lim) were performed in delft media containing 0.471 g/L of ammonium sulphate, 27.5 g/L glucose, 0.5 g/L magnesium sulphate heptahydrate, 3 g/L monopotassium phosphate, 1 mL of trace metals solution, and 1 mL of vitamin solution. Trace metal and vitamin solutions have the same composition as in shake flask cultivations. The pH was set at 5 with 2M KOH and was kept at 5 during the fermentation by automatic KOH addition.

### Extracellular metabolite analysis

Samples for high-performance liquid chromatography (HPLC) were taken after at least four residence times of steady-state cultivation. 1 mL of culture was centrifuged (5 min, 3000 rcf), and the supernatant was used for HPLC analysis. Acetate, citrate, ethanol, glycerol, glucose, pyruvate, and succinate concentrations were quantified. The HPLC system UltiMate® 3000 (Dionex) was equipped with an Aminex® HPX-87H ion exclusion column (Bio-Rad). 5 mM H_2_SO_4_ at a flow rate of 0.6 mL/min was used as eluent. Glucose was quantified using a refractive index detector (Shodex ri-101).

### Lipid extraction and quantification

Samples for lipid extraction and quantification were taken after at least four residence times of steady-state cultivation. The protocol used was previously described^23,50^. Briefly, 1 mL of cell culture was spun down (5 min, 5000 rcf), the supernatant discarded, and the pellet washed twice with 1 mL water. The resuspended cells were spun down (5 min, 5000 rcf), and the pellet resuspended in 100 μL of water. The suspension was dried in a vacuum dry freezer for 1 day. 40 μg of triheptadecanoin (TAG(17:0/17:0/17:0)) were then added to the freeze-dried cell pellet as an internal standard. After adding 500 μL of 1M NaOH in methanol the samples were vortexed at 1200 rpm for 1 h (room temperature). After adding 80 μL of 49% sulfuric acid, FAMEs were extracted by adding 500 μL hexane. Phases were separated by centrifugation (1 min, 10000 rcf) and 200 μL of the upper hexane phase were diluted 1:5 in hexane. 1 μL was analyzed on GC-MS (Thermo Scientific Trace 1310 coupled to a Thermo Scientific ISQ LT with a ZBFAME column (Phenomenex, length: 20 m; Inner Diameter: 0.18 mm; Film Thickness: 0.15 μm)). The method consisted of 2 min hold at 80°C, followed by a ramp of 40°C/min until 160°C, a ramp of 5°C/min until 185°C, a ramp of 40°C/min until 260°C and a final hold of 260°C for 30 s.

Lipid content was calculated as lipid content percentage over the cell dry weight. Cell dry weight was determined by vacuum filtration of 1 mL of samples on pre-weighed 0.45 µm filter membranes (Sartorius Biolab), followed by 15 min microwaving at 325 W and placement in a desiccator for at least 3 days.

### Solid Phase Extraction

Samples for solid phase extraction were taken after at least four residence times of steady-state cultivation. The protocol has been adapted from ^24^. Briefly, after adding 1 mL of chloroform:methanol 2:1 (v:v) (Merck), the sample was vortexed at 1200 rpm for 1 h at room temperature. SPE cartridges (Supelclean™ LC-NH2 SPE Tube, Sigma-Aldrich) were placed on Vac Elut apparatus and activated with 4 mL of hexane. Samples were then loaded on the cartridges and eluation of the different lipid fractions was achieved by adding solvents in the following order. 4 mL of chloroform:isopropanol 2:1 (v:v)(Merck) were used to elute the neutral lipid fraction (NL), including cholesterol, cholesterol ester, triacylglycerol (TAG), diacylglycerol (DAG), monoacylglycerol (MAG). 4 mL of 2% acetic acid in diethyl ether (Sigma-Aldrich) were used to elute the free fatty acid fraction. 4 mL of methanol was used to elute the phospholipid fraction. The lipid fractions were collected and the solvent was evaporated in a MiVac apparatus (35°C, 1 h). Pellets were then resuspended in 1 mL chloroform:methanol 2:1 (v:v), transferred to 1.5 mL reaction tubes, and subject to FAME extraction before analysis on GC–MS. For SPE samples the GLC reference standard GLC 426 was used, and the method consisted of 2 min hold at 80°C, followed by a ramp of 40°C/min until 120°C, a ramp of 3.5°C/min until 165°C, a ramp of 40°C/min until 260°C and a final hold of 260°C for 30 s.

### RNA-extraction and sequencing

Samples for RNA extraction were taken after at least four residence times of steady-state cultivation. 10 mL of culture were rapidly withdrawn and injected into a 50 mL falcon tube containing ca. 35 mL of crushed ice. Samples were immediately centrifuged (5 min, 4000 rcf, 4°C). After discarding the supernatant, the pellet was resuspended in 2 mL ice-cold water, transferred into 2 mL reaction tubes, and centrifuged (5 min, 4000 rcf, 4°C). The supernatant was discarded, and to remove the remaining supernatant residues, reaction tubes were tapped on paper towels. Samples were immediately frozen in liquid nitrogen and stored at −80 °C until RNA extraction.

RNA was extracted using the RNeasy Mini Kit (QIAGEN). DNA present in the samples was digested using RNase-Free DNase Set (QIAGEN). The quality of RNA samples was analyzed with a 2100 Bioanalyzer (Agilent Technologies, Inc., Santa Clara, CA). The purified RNA was stored at −80°C until further analysis.

The RNA library was constructed using Illumina TruSeq Stranded mRNA (poly-A selection) and samples were sequenced on NovaSeq6000 (NovaSeq Control Software 1.7.5/RTA v3.4.4) with a 151nt(Read1)-10nt(Index1)-10nt(Index2)-151nt(Read2) setup using ‘NovaSeqXp’ workflow in ‘S4’ mode flowcell. The Bcl to FastQ conversion was performed using bcl2fastq_v2.20.0.422 from the CASAVA software suite. The quality scale used was Sanger / phred33 / Illumina 1.8+. The raw data can be retrieved from ArrayExpress with access number E-MTAB-11008.

### Differential gene expression and gene set analysis

The raw reads were processed with the NGI RNAseq Pipeline (https://github.com/nf-core/rnaseq.git), version 3.5. The *Y. lipolytica* strain CLIB89(W29) reference genome was used to map the reads (assembly GCA_001761485.1). Differential gene expression analysis was performed with voom-limma^51,52^, and adjusted *P*-values were adjusted according to the Benjamini–Hochberg method. Volcano plots were made with the R package EnhancedVolcano^53^. OmicsBox (*https://www.biobam.com/omicsbox*) was used to generate gene sets by blasting *Y. lipolytica* exons against the RefSeq non-redundant proteins database using BlastX algorithm. A total of 31.421 GO terms were annotated to 5629 genes. Gene set analysis was performed using the R package PIANO (Platform for Integrative Analysis of Omics)^30^, using gene levels statistics, and excluding gene sets containing less than five or more than 500 genes. The code used for the analysis is available on GitHub (https://github.com/SysBioChalmers/Yarrowia_Multifactor).

## DATA AND CODE AVAILABILITY

The raw RNA-seq data are deposited in ArrayExpress with accession number E-MTAB-11008. The code and data used for the analysis are deposited on GitHub and are available at https://github.com/SysBioChalmers/Yarrowia_Multifactor.

## Acknowledgements

The authors acknowledge support from the National Genomics Infrastructure in Stockholm funded by Science for Life Laboratory, the Knut and Alice Wallenberg Foundation and the Swedish Research Council, and SNIC/Uppsala Multidisciplinary Center for Advanced Computational Science for assistance with massively parallel sequencing and access to the UPPMAX computational infrastructure.

The computations were enabled by resources provided by the Swedish National Infrastructure for Computing (SNIC) at Chalmers Centre for Computational Science and Engineering (C3SE) partially funded by the Swedish Research Council through grant agreement no. 2018-05973.

This research was funded by the Novo Nordisk Foundation [grant **NNF20CC0035580**], Research Council for Environment, Agricultural Sciences, and Spatial Planning (Formas) [grant 2018-00597], and the Swedish Research Council (VR) [grant 2019-04624].

