## Supplementary for "Abolishing storage lipids induces protein misfolding and stress responses in *Yarrowia lipolytica*"

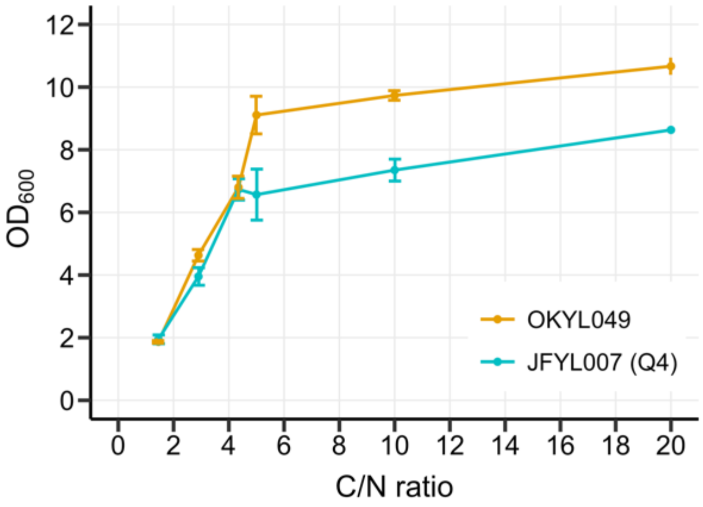


**Figure S1**: *Y. lipolytica* was grown in delft media for 72 hours. The media composition was kept constant, but the glucose concentration was varied to produce C/N ratios between 1.45 and 20. The $OD_{600}$ was measured and plotted against the C/N ratio. We tested the Q4 strain (JFYL07) and an obese strain (OKYL049) as they show opposite phenotypes that might affect the threshold between carbon or nitrogen limitation. Q4 lacks the genes responsible for TAGs synthesis. OKYL049 (*DGA1* overexpression and *are1* deletion) accumulates high levels of TAGs. C/N ratios between 1.45 and 4.43 are carbon limiting for both strains. At higher C/N ratios, the nitrogen becomes limiting, and increasing the glucose concentration doesn't have a major effect on the final $OD_{600}$. Dots represent the average $OD_{600}$ of triplicates, and error bars represent the standard deviation.

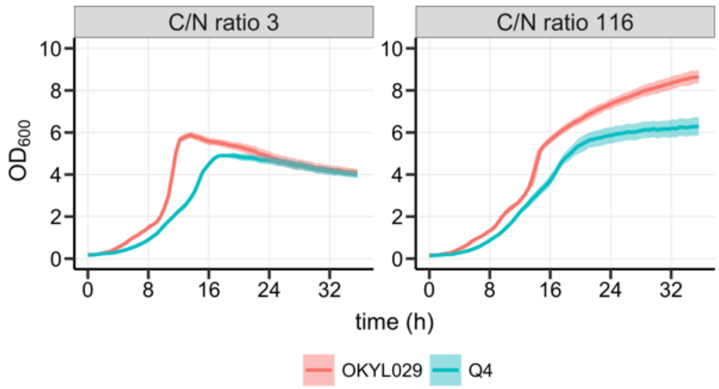


**Figure S2**: Strains OKYL029 and Q4 were cultured in 96-wells plates with C/N ratio 3 (C-lim, left panels) or C/N ratio 116 (N-lim, right panels). $OD_{600}$ was measured with the growth profiler every 30 minutes. The curves represent the average of triplicates, and the shadowed areas represent the standard deviation.

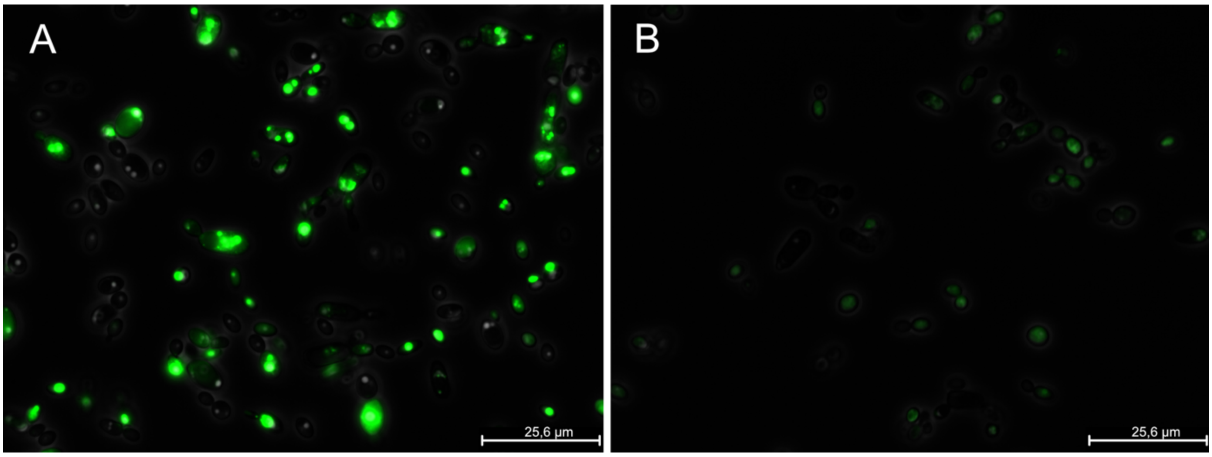


**Figure S3:** Microscope pictures of *Y. lipolytica* strain OKYL029 (A) and Q4 (B) grown under N-lim. Cells were cultured in N-lim Novogy media (0.5 g/L urea, 1.5 g/L yeast extract, 0.85 g/L casamino acids, 1.7 g/L yeast nitrogen base (YNB) without amino acids and ammonium sulphate, 5.1 g/L potassium hydrogen phthalate, 100.0 g/L glucose, pH set to 5.5 with KOH) for 72 hours and stained with Bodipy® Lipid Probe. In the wild-type strain OKYL029 (A) lipid droplets (LD) are visible. No visible lipid droplets (LD) were observed in the Q4 strain.

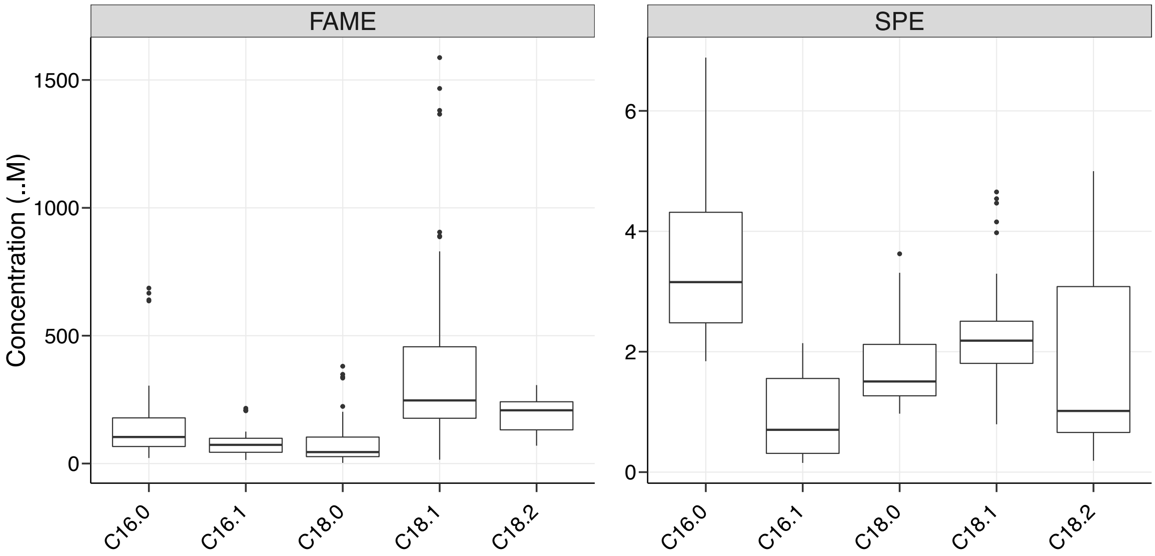


**Figure S4:** Molar concentrations of each fatty acid (C16:0, C16:1, C18:0, C18:1, C18:2) both in total lipids and in the FFA fraction. The concentrations of lipids have been calculated based on the FAME and SPE data. The highest concentration measured is 1500 µM for the total lipids, while for the FFA fraction is 6 µM.
